# A tessellated lymphoid network provides whole-body T cell surveillance in zebrafish

**DOI:** 10.1101/2023.01.17.524414

**Authors:** Tanner F. Robertson, Yiran Hou, Simone Shen, Julie Rindy, John-Demian Sauer, Huy Q Dinh, Anna Huttenlocher

**Author notes:** Corresponding author: Anna Huttenlocher. **Author Contributions:** Conceptualization: T.F.R, A.H. Methodology: T.F.R., S.S., J.D.S, A.H. Investigation: T.F.R. Visualization: T.F.R, Y.H. Funding acquisition: T.R., A.H. RNA-sequencing analysis: Y.H., H.Q.D. Supervision: J.D.S, H.Q.D, A.H. Project Administration: T.F.R, J.R., A.H. Writing – original draft: T.F.R. Writing – review and editing: T.F.R, Y.H., S.S, J.D.S., H.Q.D, A.H. **Competing Interest Statement:** The authors declare that they have no competing interests.

## Abstract

Homeostatic trafficking to lymph nodes allows T cells to efficiently survey the host for cognate antigen. Non-mammalian jawed vertebrates lack lymph nodes but maintain similarly diverse T cell pools. Here, we exploit *in vivo* imaging of transparent zebrafish to investigate how T cells organize and survey for antigen in an animal devoid of lymph nodes. We find that naïve-like T cells in zebrafish organize into a previously undescribed whole-body lymphoid network that supports streaming migration and coordinated trafficking through the host. This network has the cellular hallmarks of a mammalian lymph node, including naïve T cells and CCR7-ligand expressing non-hematopoietic cells, and facilitates rapid collective migration. During infection, T cells transition to a random walk that supports antigen presenting cell interactions and subsequent activation. Our results reveal that T cells can toggle between collective migration and individual random walks to prioritize either large-scale trafficking or antigen search *in situ*. This novel lymphoid network thus facilitates whole-body T cell trafficking and antigen surveillance in the absence of a lymph node system.

**Significance Statement:** In mammals, lymph nodes play a critical role in the initiation of adaptive immune responses by providing a dedicated place for T cells to scan antigen-presenting cells. Birds, reptiles, amphibians, and fish all maintain diverse repertoires of T cells but lack lymph nodes, raising questions about how adaptive immunity functions in lower jawed vertebrates. Here, we describe a novel network of lymphocytes in zebrafish that supports whole-body T cell trafficking and provides a site for antigen search, mirroring the function of mammalian lymph nodes. Within this network, T cells can prioritize large-scale trafficking or antigen scanning by toggling between two distinct modes of migration. This network provides valuable insights into the evolution of adaptive immunity.

## Introduction

Adaptive immunity relies on rare, naïve T cells finding cognate antigen that arise in the body’s many diverse tissues during an infection (1). In mammals, antigen discovery and subsequent T cell activation occur primarily in lymph nodes, regional organs that provide a specialized space for T cells to efficiently migrate and scan antigen-presenting cells (APCs) (1, 2). In this lymph node system, naïve T cells serially enter, search within, and exit lymph nodes that are distributed strategically throughout the host, allowing individual T cell clones to perform whole-body antigen surveillance on the timescale of days (1, 3).

While lymph nodes are largely restricted to mammals, lymphocytes with diverse antigen receptors are present in all jawed vertebrates (4, 5). It remains unclear how T cells in fish, reptiles, amphibians, and birds effectively survey for antigen in a host devoid of lymph nodes. Further, since diverse lymphocyte repertoires emerged approximately 500 million years before lymph nodes, adaptive immunity necessarily evolved a different surveillance strategy than the one operating in mammals (4, 5). While lymphocytes typically populate and even form lymphoid tissues around the respiratory and digestive organs of lower jawed vertebrates (6, 7), these animals produce robust adaptive immune responses when these barrier sites are bypassed by intraperitoneal or intramuscular immunization (8, 9). These data indicate that non-mammalian jawed vertebrates also perform whole-body antigen surveillance, but how T cells effectively search for antigen in the absence of lymph nodes remains unknown.

Studies in transparent zebrafish have previously uncovered new mechanisms of immune cell migration and behavior *in situ (10-13)*, and this model organism provides a unique opportunity to investigate T cell surveillance in an organism lacking lymph nodes. Utilizing a combination of microscopy and single cell RNA-sequencing (scRNA-seq), we first identified a novel whole-body lymphoid tissue that encompasses the major cellular hallmarks of a mammalian lymph node. Live imaging revealed that T cell can toggle between two major modes of motility within this novel lymphoid tissue: one that uses T cell multicellular streaming to facilitate long-distance coordinated movement and a second more individual random walk that facilitates antigen-presenting (APC) cell scanning. We find that infection drives a switch from streaming to a random walk and provide evidence that T cells are detecting antigen, activating, and differentiating within this lymphoid network. Collectively, we show that zebrafish T cells utilize distinct modes of motility within this novel lymphoid network to perform whole-body antigen surveillance, all in the absence of lymph nodes.

## Results

### T cells organize into a tessellated lymphoid network in juvenile and adult zebrafish

Prior studies have used larval stage animals to study T cell motility, however a functional adaptive immune system does not emerge until 4-6 weeks post-fertilization (wpf) (14, 15). We therefore investigated T cell distribution and trafficking in 10 wpf transparent zebrafish when the adaptive immune system is well-established (15-17). Organism-scale imaging of tg*(lck:GFP)* revealed that T cells organize into a regularly repeating hexagonal pattern across the body of the animal (Fig. 1A). This hexagonal pattern was unique to the body, with T cells in the fins and head assuming a more diffuse distribution (Fig. 1A, S1A-B). Within this hexagonal pattern, GFP-labelled cells were particularly abundant at the dorsal and ventral poles, giving the network a zigzag appearance (Fig. 1A). This pattern is completely absent in the more well-studied larval zebrafish but becomes progressively apparent throughout the juvenile stages of development and is maintained into adulthood (Fig. 1B). Individual T cells are apparent in larval fish (Fig. S1C), but they do not organize into this higher order pattern (Fig. 1B). Notably, the emergence of this T cell pattern corresponds temporally with the maturation of adaptive immunity (15).

**Figure 1.**
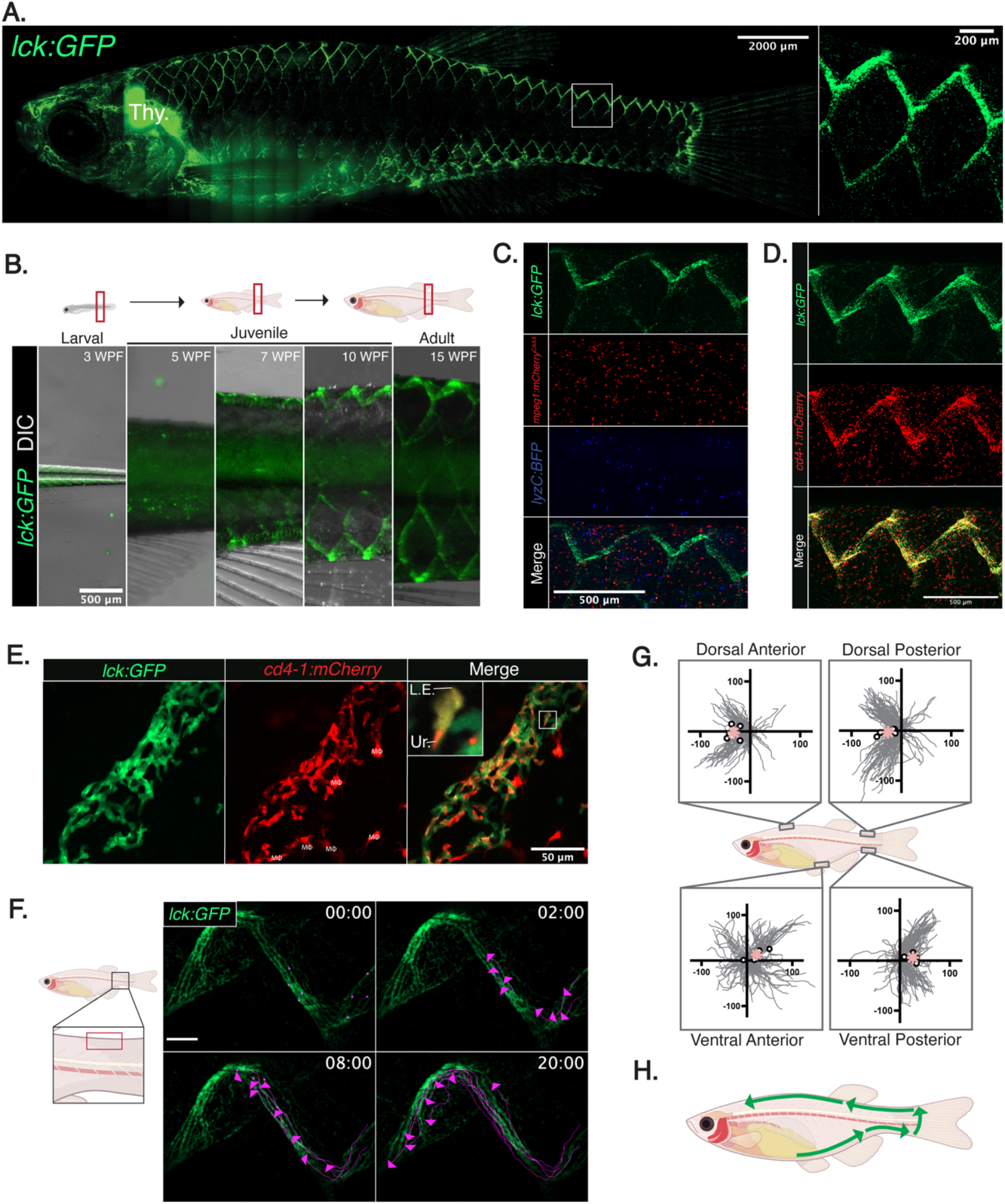
A tessellated lymphoid network supports coordinated whole-body T cell trafficking in juvenile and adult zebrafish. **(A)** The organism-scale organization of T cells in a 10 wpf tg(lck:GFP) zebrafish with the inset showing the hexagonal ‘zigzag’ pattern. **(B)** The emergence of the T cell network is shown with representative images of the caudal peduncle of tg(lck:GFP) zebrafish at 3, 5, 7, 10, and 15 wpf. **(C)** Imaging of the caudal peduncle shows the organization of T cells (lck:GFP), macrophages (mpeg1:mCherry^CAAX^), and neutrophils (lyzc:BFP) in triple transgenic zebrafish. **(D-E)** Imaging of the caudal peduncle at low (d) or high (e) magnification shows the presence of dual labelled cells in tg(lck:GFP;cd4-1:mCherry) fish. MΦ = macrophage, L.E. = leading edge, Ur. = uropod. **(F)** Timelapse imaging of T cells migrating through the TLN over 20 minutes, with the cell tracks and directions of 8 representative GFP+ cells shown in magenta. Scale bar = 100 microns. **(G)** Plots show the cell tracks of GFP+ cells migrating in the TLN of tg(lck:GFP;cd4-1:mCherry) in the indicated regions of 5 individual fish pooled from 3 independent experiments. Each dot = average per fish, asterisks = average total movement. Scale is in microns. **(H)** A summary of the movement of T cells in the TLN.

To our knowledge, a network that organizes T cells into a repeating pattern has never been observed in any organism. To determine if other leukocyte populations assume a similar distribution, we generated triple transgenic *lck:GFP;mpeg1:mCherry*^*CAAX*^*;lyzc:BFP* zebrafish to visualize T cells, macrophages, and neutrophils simultaneously (17-19). These animals revealed that this zigzag pattern was unique to T cells, with both macrophages and neutrophils showing a more diffuse organization (Fig. 1C). Using double transgenic *lck:GFP;cd4-1:mCherry* fish (20), we found mCherry-expressing T cells to be common within the TLN (Fig. 1D-E) but infrequent in larval zebrafish (Fig. S1C). GFP-negative, mCherry-expressing macrophages, described previously (20), are also apparent in proximity to the TLN (Fig. 1E). Reflecting its repeating pattern, we named this network the Tessellated Lymphoid Network (TLN).

### T cells traffic through the host in a coordinated ventral-to-dorsal loop

TLN T cells commonly assume the “hand-mirror” morphology typical of motile lymphocytes, prompting us to perform live imaging experiments (Fig. 1E, inset). TLN T cells migrated in a remarkably coordinated and directional manner, with cells maintaining directional persistence over hundreds of microns as they zigzagged through the network (Fig. 1F, Movie S1). To understand this movement on the organism-scale, we performed live imaging in four discrete positions: dorsal anterior, dorsal posterior, ventral anterior, and ventral posterior (Fig. 1G). Remarkably, T cells showed clear rostral movement on the dorsal side and clear caudal movement on the ventral side (Fig. 1G-H). The TLN is punctuated on the rostral and caudal ends of the body by large reservoirs of T cells (Fig. S1D-E). Live imaging is technically challenging at the rostral reservoir due to the presence of the operculum and pectoral fin, but within the caudal reservoir we observed clear T cells movement from the ventral TLN to the dorsal TLN (Fig. S1F-I). These data collectively suggest that the ventral and dorsal portions of the TLN are continuous, allowing T cells to traffic through the host in a coordinated ventral-to-dorsal loop (Fig. 1H).

### The TLN localizes to a pocket formed by overlapping scales and is the largest reservoir of T cells in zebrafish

Fish scales organize in a hexagonal pattern, like the TLN (21). To determine if the TLN associates with fish scales, we imaged adult tg(*lck:GFP*) zebrafish stained with alizarin red and found the TLN and scale pattern to be clearly interrelated (Fig. 2A). Single z-section images showed that the TLN localizes to narrow regions formed between adjacent scales (Fig 2B), positioning this network just underneath the barrier surface of the animal (Fig. 2C). Given this localization, we next investigated if TLN T cells could be isolated by descaling adult zebrafish (Fig. 2D-E). Descaling resulted in an almost complete loss of T cells from the TLN without causing loss of T cells from the nearby fins (Fig. 2D-E). When descaling was performed on adult tg(*lck:GFP*) zebrafish in isotonic buffer, GFP-expressing cells from the TLN appeared as a suspension with GFP-negative cells (Fig. 2F), and flow cytometric analysis of this suspension revealed that the GFP-positive cells were live and localized to the lymphocyte gate as expected (Fig. S2A-B). We characterized the TLN by flow cytometry and compared it to other known reservoirs of T cells in fish: the gills, gut, kidney, and spleen (Fig. 2G) (6, 7, 20). The TLN, gills, and gut all had a similar proportion of GFP-positive cells (Fig. 2G), but the TLN harbored more total live cells than all sites tested besides the kidney (Fig. 2H). In total, we harvested nearly 1×10^5^ GFP-positive cells from the TLN of each 15 wpf tg(*lck:GFP*) fish, more than all other sites tested (Fig. 2I). Using tg(*lck:GFP*;*cd4-1:mCherry*) zebrafish, we found that approximately 66% of the TLN T cells also expressed *cd4-1* (Fig. S2C). Other groups have estimated that adult zebrafish have 2×10^5^ total T cells (22), suggesting that this lymphoid network is the largest reservoir of T cells in the host.

**Figure 2.**
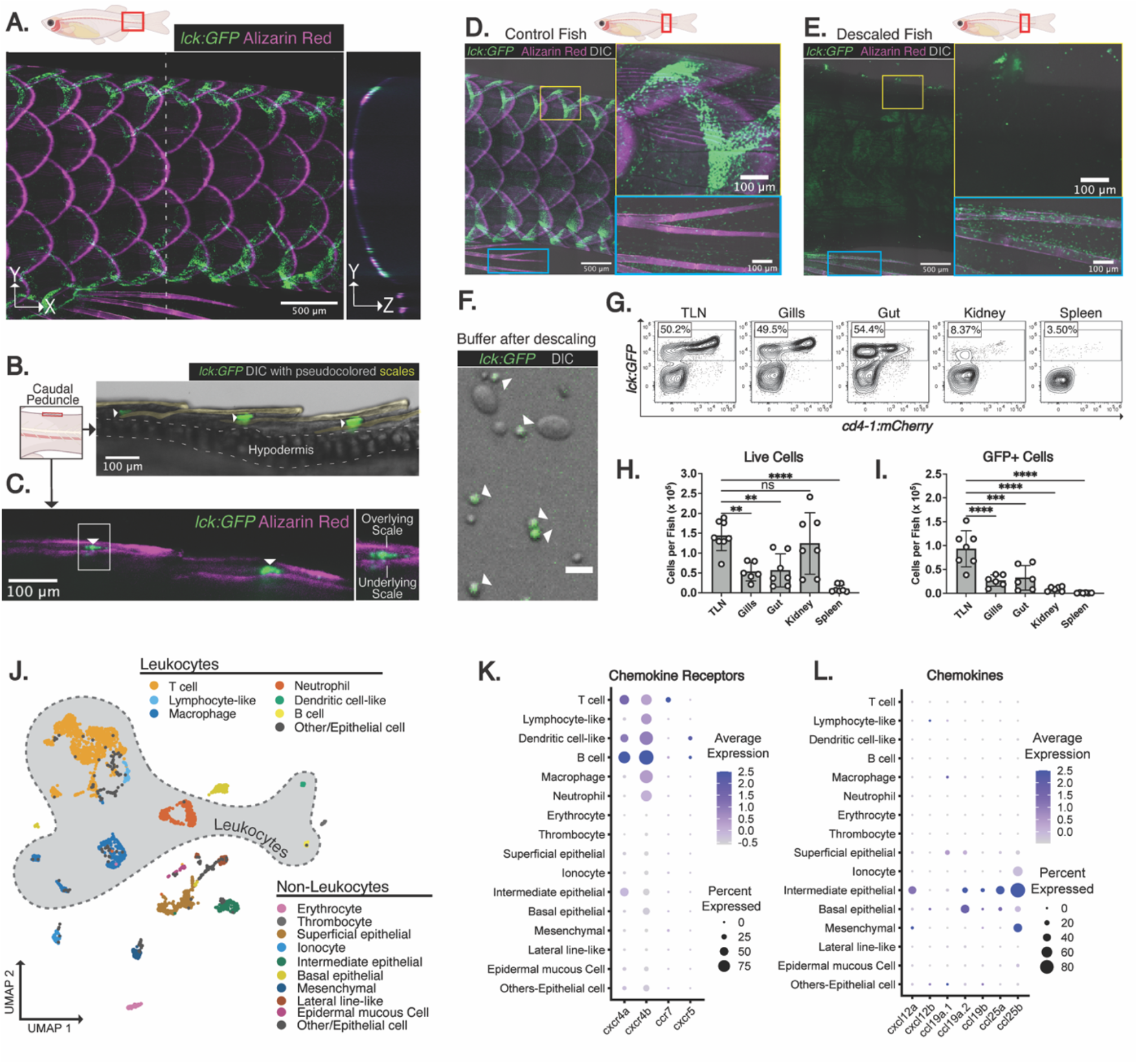
The TLN is the largest reservoir of zebrafish T cells and shares the cellular hallmarks of a mammalian lymph node. **(A)** The relationship between the T cell pattern and scales shown by imaging the caudal peduncle of tg(lck:GFP) fish stained with Alizarin red. **(B)** The tissue localization of the T cell network shown by imaging a single *z*-plane of the caudal peduncle of a tg(lck:GFP) fish, with the yellow transparent overlays emphasizing the scales. **(C)** Imaging of a tg(lck:GFP) zebrafish stained with Alizarin red shows T cells (white arrows) in a pocket formed by overlapping scales. **(D-E)** Imaging of control and descaled tg(lck:GFP) adult zebrafish stained with alizarin shows the loss of the TLN T cells upon descaling (yellow inset) but not fin T cells (blue inset). **(F)** After descaling tg(lck:GFP) zebrafish, the buffer contains a single-cell suspension of GFP-expressing (arrows) and GFP-negative cells. Scale bar = 10 microns. **(G-I)** Representative flow plots (G) and enumeration of live (H) and GFP-expressing (I) cells harvested from the indicated organ of tg(lck:GFP/cd4.1:mCherry) 15 wpf zebrafish, pre-gated on live singlets, with the frequency of GFP-expressing cells shown. (E) Each dot corresponds to a single fish showing 5-7 individuals pooled from four separate experiments. The mean of the TLN was compared to the mean of every other group with an ordinary one-way ANOVA. **(J)** RNA-seq profiling for all cells collected by descaling adult zebrafish, with leukocyte populations outlined in gray. **(K-L)** Representative gene expression in each of the major cell types as a dot plot showing select chemokine receptor (K) and chemokine (L) expression.

### The TLN harbors naïve-like T cells, APCs, and chemokine-producing non-hematopoietic cells

Three distinct cellular populations work in synchrony to facilitate antigen surveillance in mammalian lymph nodes: naïve T cells, antigen-presenting cells, and chemokine-producing fibroblastic reticular cells (FRCs) (2, 23-25). To determine if these cell populations exist within the TLN, we performed scRNA-seq on the cells harvested by descaling. Leukocytes made up the majority of identified cells, with T cells making up the most abundant cell type (Fig. 2J, Fig. S2D). Based on the expression pattern of lineage-defining genes, we also identified all three major APC populations (macrophages, dendritic-like cells, B cells) as well as a variety of other leukocytes (Fig. 2J, Fig. S3E). Further clustering of the T cells revealed subpopulations, including two large clusters of naïve-like T cells that expressed homeostatic chemokine receptors (*ccr7, cxcr4a*) but showed little to no expression of various cytokines (*il4, il13, il11b*), effector proteins (*gzma, nkl1*.*2*), or markers of proliferation (*mki67, top2a*) (Fig. S2I-J). We also identified a regulatory *foxp3a-*expressing T cell population, a small population of T cells exhibiting constitutive expression of type-2 cytokines (*il4, il13, il11b*; Th2/ILC2 cell), two discrete populations with gene signatures corresponding to cytotoxic T cells (Cytotoxic T cell-1 and 2), a population expressing proliferation markers (*mki67, top2a*; Cycling T cell), and various other minor T cell populations (Fig. S2I-J).

T cell and APC organization and migration within lymph nodes is chiefly dictated by the specific expression of ligands for CXCR4 (CXCL12), CCR7 (CCL19, CCL21) and CXCR5 (CXCL13) by different FRC subsets (25). Because of gene duplication events, zebrafish maintain two orthologues of CXCL12 (*cxcl12a, cxcl12b*), three orthologues of CCL19 (*ccl19a*.*1, ccl19a*.*2, ccl19b*), two genes that are putative orthologues of both CCL21 and CCL25 (*ccl25a, ccl25b*), and one orthologue of CXCL13 (*cxcl13*) (26-28). The chemokine receptor expression profile for leukocytes in the TLN was similar to lymph nodes (25). All leukocytes expressed at least one of the two CXCR4 orthologues (*cxcr4a, cxcrb*) (Fig. 2K). Expression of *ccr7* was only appreciably detected in T cells, and this expression was concentrated in the naïve-like populations (Fig. 2K, Fig. S2L). Conversely, *cxcr5* was detected in a minority of dendritic-like cells and B cells (Fig. 2K). We next sought to identify the cell population producing the corresponding chemokines for these receptors. As expected, expression of the relevant chemokines was virtually undetectable in all the hematopoietic cell populations (Fig. 2L). Intermediate epithelial cells, a poorly understood population of cells in the fish skin, expressed high levels of *cxcl12a* as well as all the known CCR7 ligands except for *ccl19a*.*1* (Fig. 2L). Although basal epithelial cells and mesenchymal cells also showed appreciable expression levels of *ccl19a*.*2* and *ccl25b* respectively, intermediate epithelial cells appeared to be the primary source of CCR7 ligands (Fig. 2L). Notably, leukocytes that lack *ccr7* expression (macrophages, neutrophils) showed a diffuse rather than tessellated distribution, supporting a role for this chemokine ligand-receptor pairs in organizing the TLN (Fig. 1C).

### TLN T cells use ballistic multicellular streaming to migrate

The TLN maintains all the cellular components that permit effective antigen surveillance in mammalian lymph nodes, so we investigated TLN T cell search strategy and migration using real time imaging with tg(*lck:GFP;cd4-1:mCherry*) fish. Far from the random walk that typifies migration in lymph nodes, TLN T cells moved in a remarkably coordinated manner, with nearly every T cell moving in the same direction (Fig. 3A, Movie S2). TLN T cells migrated at an average velocity of 14.2 microns/min and with consistently high directionality (Fig. 3B-C). Unlike in a random walk, where mean-squared displacement (MSD) increases linearly with time (2, 29-32), cell tracking revealed that TLN T cells instead showed a superdiffusive mode of motility that correlates linearly with displacement rather than MSD (Fig. 3D-E). The coordinated nature of this movement can be explained by the fact that T cells in the TLN form frequent and prominent head-to-tail interactions, as observed between the leading edge and uropod of mCherry-positive and mCherry-negative T cells in tg(*lck:GFP*;*cd4-1:mCherry*) fish (Fig. 3F). In younger juvenile fish where the TLN is sparsely populated, T cells can be seen forming, breaking, and re-forming these associations multiple times as they migrate in multicellular units (Fig. 3G). As the TLN matures, these interactions become stable and T cells form a continuum that resembles a flowing fibrous network (Fig. 3H-I, Movie S3). This stable multicellular streaming resembles dictyostelium discoideum streaming motility (33) and explains how T cells maintain directional persistence over hundreds of microns within the TLN (Fig. 1F, Movie S1).

**Figure 3.**
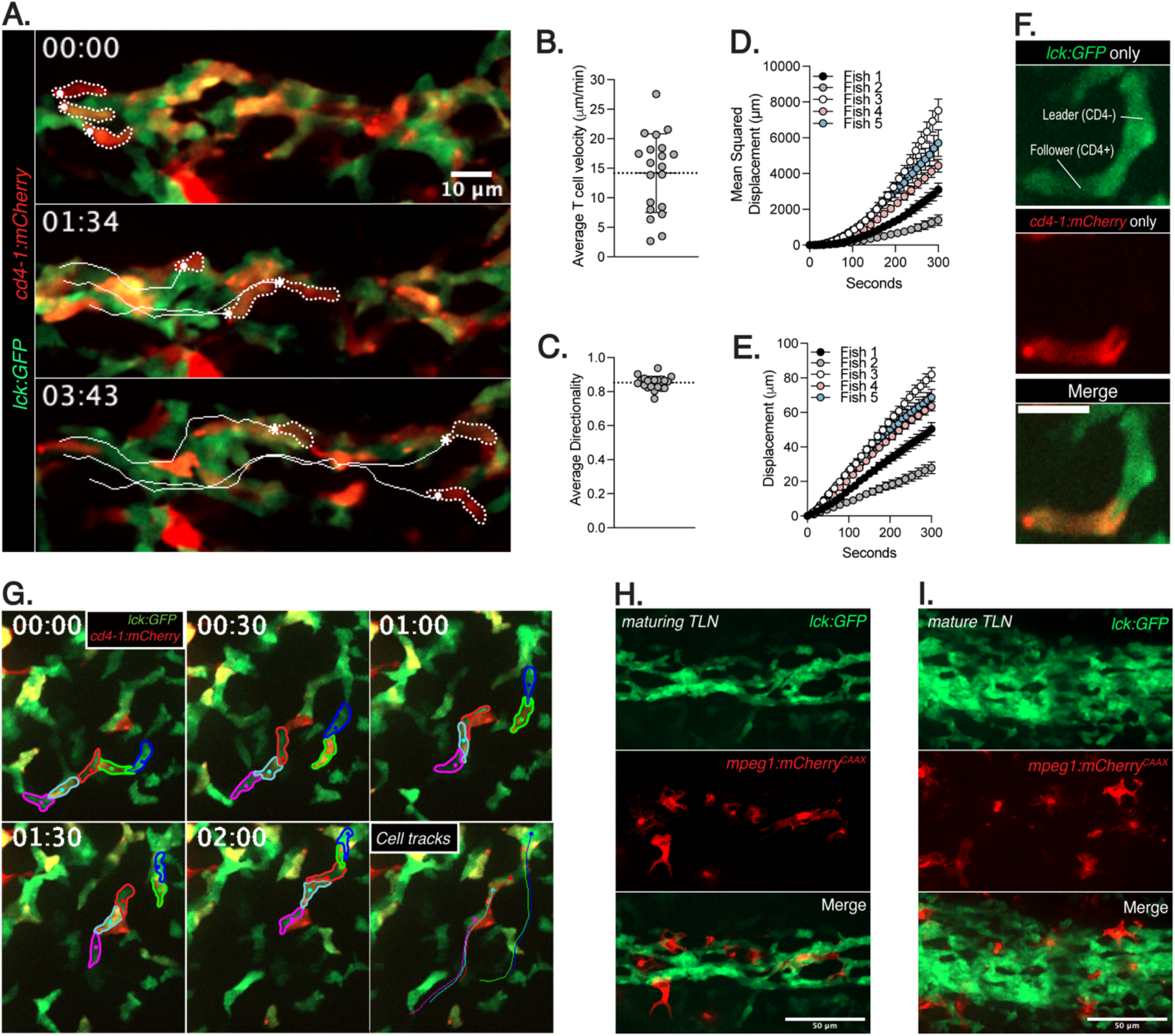
TLN T cells use ballistic multicellular streaming to migrate. **(A)** Timelapse imaging of the TLN in adult tg(lck:GFP/cd4-1:mCherry) zebrafish shows the directional migration of 3 outlined T cells. **(B-C)** Average T cell velocity (B) and directionality (C) of T cells within the TLN, with each dot corresponding to the average of all T cells from an individual fish pooled from three independent experiments. **(D-E)** Plots show the average mean squared displacement (D) and average displacement (E) vs time of T cells migrating in the TLN. **(F)** Imaging shows the head-to-tail interaction between two T cells (B) mm:ss, scale = 10 microns. **(G)** Timelapse imaging shows five outlined T cells in a tg(lck:GFP/cd4-1:mCherry) zebrafish forming, breaking, and re-forming head-to-tail interactions as they migrate through the TLN of a 6 WPF juvenile fish, with the final frame showing the cell tracks. **(H-I)** Imaging of the TLN in tg(lck:GFP/mpeg1:mCherry) shows T cells organizing into a fibrous network with embedded macrophages as TLN goes from maturing (H) to fully mature (I) in late juvenile or adult zebrafish.

### A streaming to random walk motility switch occurs during infection

While streaming motility is well-suited for trafficking through the host, it appears ill-suited for antigen search. Macrophages are stably embedded within the TLN, but streaming T cells contact these APCs only transiently (Fig. 3H-I, Movie S3). We hypothesized that this behavior may change during infections. To this end, we developed a localized bacterial infection model by puncturing the dorsal side of the caudal peduncle with a needle and incubating the wounded fish for 2 hours in a bath of mCherry-expressing *L. monocytogenes* (Fig. S3A). The infection expands and resolves over a 7-day timespan in most fish, with TLN T cells accumulating in regions proximal to the infected wound between 1-5 days post-infection (dpi; Fig. S3B-C, Fig. 4A-B).

**Figure 4.**
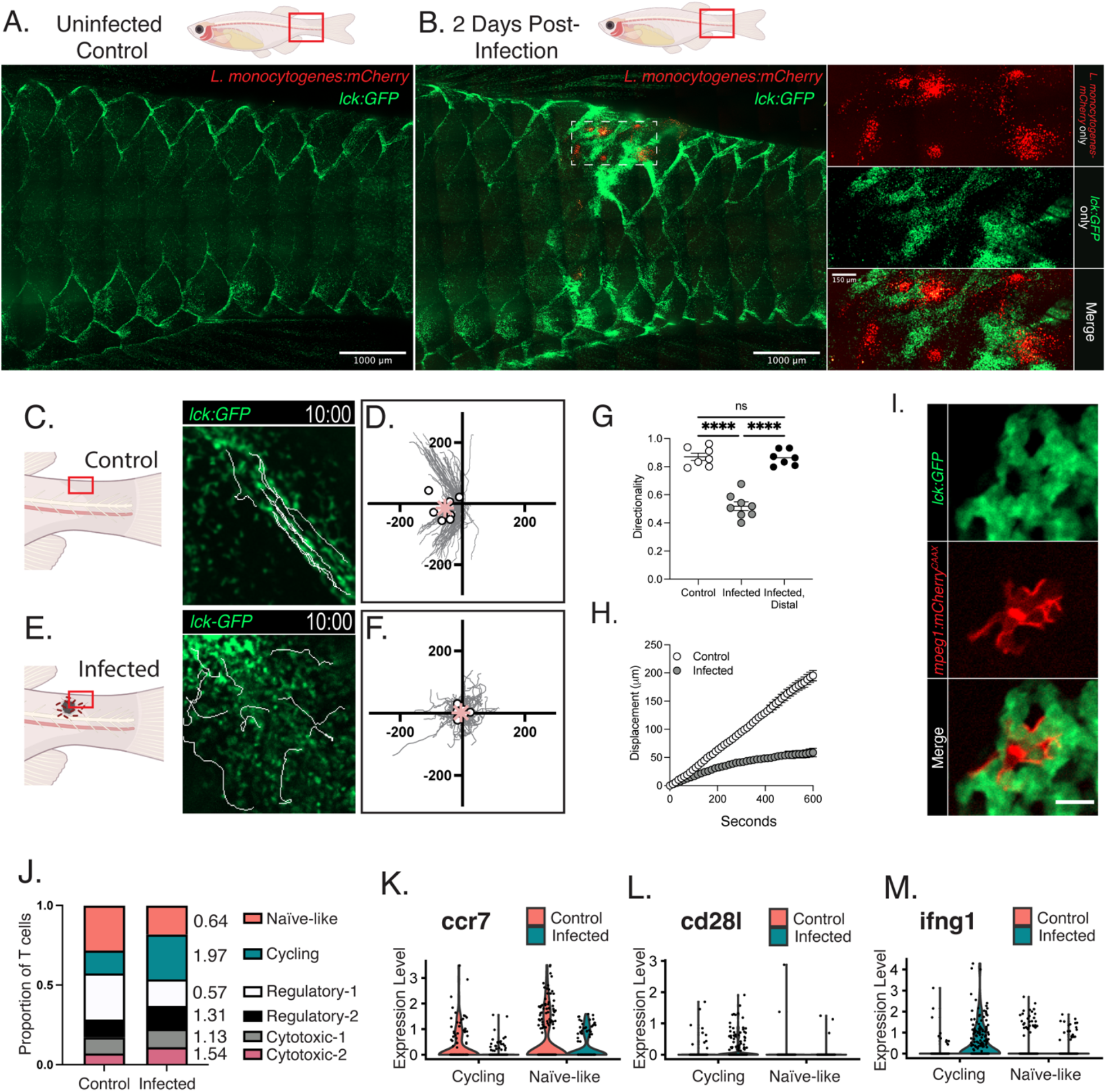
TLN T cells transition to a random walk during infection to scan APCs. **(A-B)** T cells accumulate around infections in tg(lck:GFP) zebrafish at 2 dpi with mCherry-expressing *L. monocytogenes*. **(C, E)** Altered T cell motility at 2 dpi (E) compared to control (C), with 10 overlaid cell tracks **(D, F)** 10-minute cell tracks from 7-8 uninfected (d) or 2 dpi (f) *tg(lck:GFP;cd4-1:mCherry)* fish pooled from 3 independent experiments. Dots = average movement per fish, asterisk = average total movement. The scale is in microns. **(G)** Infected-proximal but not infected-distal T cells exhibit reduced directionality, with each dot corresponding to the average of an individual fish pooled from 3 independent experiments. Means compared with an ordinary one-way ANOVA. **(H)** Representative displacement vs time plots show the loss of superdiffusive motility in infection-proximal T cells in fish at 2 dpi. **(I)** T cells associate with infection proximal macrophages in tg(lck:GFP/mpeg1.1-mcherry^CAAX^) fish. Scale bar = 10 microns **(J)** Proportions of T cell subtypes in control and infected conditions with fold-change noted. **(K-M)** Violin plots show the expression levels of ccr7 (K), cd28l (L), and ifng1 (M), in cycling and naïve-like T cells harvested by descaling control or infected fish.

While TLN T cells in control fish are highly motile, T cells around the infected wound showed both motile and sessile behaviors (Movie S4). Motile T cells proximal to the infection did not exhibit ballistic streaming and instead migrated individually in apparently random directions (Fig. 4C-F, Movie S5). These infection-proximal T cells showed a sharp reduction in directionality and a sublinear relationship between time and displacement, consistent with a random walk (Fig. 4G-H). This effect was limited to infection-proximal T cells, as TLN T cells distal to the infection (>500 microns) maintained a highly directional mode of migration (Fig. 4G). Infection-proximal T cells often interacted sequentially with multiple macrophages over the course of a short random walk, similar to how T cells search for antigen in mammalian lymph nodes (Movie S6)(2, 34).

### T cell activation and differentiation within the TLN during infections

We noticed that the sessile population of T cells proximal to the infection were often in intimate, stable contact with regional macrophages (Fig. 4I). Signaling through the TCR acts as a migratory stop signal, which facilitates a stable interaction with APCs that is important for T cell proliferation and differentiation (35). To determine if TLN T cells were activated, we performed single cell analysis of the TLN harvested from infected fish. We observed several transcriptional changes consistent with T cell activation after infection. In our integrated analysis, T cells segregated into 6 discrete subsets: a naïve-like population, two *foxp3a*-expressing regulatory T cell populations, two cytotoxic T cell populations, and a cycling T cell population enriched for proliferation markers (*mki67, top2a*; Fig. S3D-E, Fig. 4J). Consistent with T cells becoming activated within the TLN, the proportion of naïve-like T cells decreased while the cycling T cells and cytotoxic T cell-2 population increased (Fig. 4J). Mirroring the TCR-dependent transcriptional changes found in mammalian T cells, *ccr7* expression was decreased and *cd28l* (orthologue of CTLA-4) was increased in T cells harvested from infected fish, especially within the cycling population (Fig. 4K-L, Fig. S3E) (36, 37). In mice, *L. monocytogenes* is a Th1-skewing infection, with activated T cells upregulating type 1 effector genes like *ifng (38-40)*. Likewise, cycling T cells harvested from infected fish had higher expression of *ifng* compared to cycling T cells from control fish or naïve-like T cells from either group (Fig. 4M, Fig. S3E). These data indicate that T cells are both activating and differentiating within the TLN during infection.

## Discussion

In summary, we have identified a novel, regularly repeating pattern of T cells in zebrafish that we have termed the Tessellated Lymphoid Network. Akin to mammalian lymph nodes, the TLN harbors *ccr7*-expressing naïve-like T cells, antigen-presenting cells, and a non-hematopoietic cell population producing CCR7 chemokine ligands. Within the TLN, T cells use a ballistic streaming mode of motility that allows them to traffic through the TLN in an efficient and coordinated ventral-to-dorsal loop. In the context of infection, TLN T cells transition to a diffusive random walk that facilitates interactions with infection-proximal APCs. Finally, scRNA-seq data provides evidence that T cells are detecting antigen, activating, and differentiating within this network. The TLN thus utilizes a different approach to solve the same problem addressed by the lymph node system in mammals: antigen may arise in distant tissues during an infection, and adaptive immune responses rely upon antigen-specific T cells finding it. By toggling between two modes of migration, ballistic streaming or a random walk, TLN T cells can prioritize either large-scale trafficking or local antigen search.

The degree to which multicellular streaming may occur in mammalian lymph nodes is unclear, but it is supported by modeling and direct observation (41). Streaming is difficult to study with intravital imaging in mammals because only a small fraction of T cells are labelled in these experiments. However, several reports have demonstrated that lymphocytes clearly possess the ability to alternate between random walks and ballistic migration. Double positive thymocytes in the outer cortex of the thymus utilize a random walk but transition into rapid ballistic migration following positive selection to move into the medulla (42), naïve lymph node B cells utilize a random walk within follicles, but migrate directionally towards the B/T cell zone boundary following antigen detection (43), and germinal center B cells employ directional migration to move between light and dark zones (44). The emerging consensus between this report and previous literature is that random walks are useful for scanning for antigen, and ballistic migration is well-suited for traveling long distances within a tissue. Ballistic streaming may be more common in the TLN than lymph nodes because internodal travel is facilitated by the blood and lymphatic vasculature, whereas TLN T cells appear to manually migrate through the host. Ballistic motility may facilitate movement to or from the blood and lymphatic vessels that facilitate lymph node entry and exit and the lymph node parenchyma, but this concept will need to be tested experimentally and remains challenging due to the lack of transparency and cellular density of mammalian lymph nodes.

The widespread use of ballistic streaming under homeostatic conditions indicates that TLN T cells are not thoroughly searching for antigen unless there is active inflammation. Lymph node T cells also alter their search strategy depending on inflammation, with some T cells exiting uninflamed lymph nodes within minutes of entering, searching for antigen minimally if at all (45). Grigorova et al. (45) have proposed a hierarchical model of antigen surveillance in lymph nodes where T cells initially survey for regional inflammation and then secondarily search for antigen. This model appears to also describe antigen surveillance in the TLN. Within this framework, streaming may help limit T cell reactivity to self-antigens by restricting antigen surveillance to scenarios where cognate antigen is likely to be found. This model also suggests that T cell search in most settings is deliberately sub-optimal and warrants careful comparisons of T cell motility in control and inflamed settings.

It will be interesting to determine how far a TLN-like system for antigen surveillance extends throughout jawed vertebrates. We favor a model where the TLN gradually evolves over time and represents the precursor to the modern mammalian lymph node. To this end, a better understanding of the molecular and genetic regulators of TLN formation will facilitate phylogenetic analysis across species. We anticipate that a deeper understanding of the evolution of adaptive immunity will provide unexpected insights into human immunology. Indeed, the zebrafish tessellated lymphoid network provides a way to study T cell antigen search without surgical manipulation, two-photon microscopy, or having to limit analysis to a small fraction of T cells. The TLN may also be useful for studying tertiary lymphoid organs in mammals, as both the TLN and TLOs are essentially unencapsulated lymph nodes (46). From a cell biology perspective, this report provides the first unambiguous evidence that lymphocytes can stream *in vivo*, providing a model to further investigate this behavior.

## Materials and Methods

### Zebrafish husbandry and maintenance

All protocols using zebrafish in this study were approved by the University of Wisconsin-Madison Research Animals Resource Center (protocol M005405-A02). Adult AB strain fish and transgenic zebrafish lines including the *Casper* WT line (16) and previously published transgenic reporter lines *lck:GFP* (17), *cd4*.*1:mCherry* (20), *mpeg1*.*1:mCherry*^*CAAX*^ (18), and *lyzC:BFP* (19) were used in this study. Double or triple transgenic reporters were generated by crossing individual lines and screening for fluorescence. For most experiments, these fish were then in-crossed to generate fish utilized for experiments. After breeding, fertilized embryos were placed into E3 media (5 mM NaCl, 0.17 mM KCl, 0.44 mM CaCl_2_, 0.33 mM MgSO_4_, 0.025 mM NaOH, and 0.0003% Methylene Blue) and maintained at 28.5°C in a petri dish in a laboratory incubator. *Lck:GFP* and *cd4-1:mCherry* larvae were screened at 6-7 days post fertilization (dpf) for fluorescence signal emanating from the paired thymus in the head. *mpeg1*.*1:mCherry*^*CAAX*^and *lyzC:BFP* reporter fish were screened looking for fluorescent macrophages and neutrophils in the caudal hematopoietic tissue anytime from 3-7 (dpf). At 7 dpf, screened larvae were transferred to a fish tank with still water where they were fed twice daily and maintained on a 14-h/10-h light/dark schedule. At 10 dpf, a drip of fresh water was applied to the tank, and at 21 dpf a gentle stream of water was applied. Fish were maintained at density of approximately 20 fish per 3 liter tank to maintain experiment-by-experiment growth consistency. Fish were removed from this environment as needed for experiments. When taking whole-animal confocal tiling images, fish were fasted overnight to reduce non-specific fluorescence coming from the intestinal tract.

### Fish Handling for Imaging

For live imaging experiments, juvenile or adult zebrafish were added to water containing 0.08 mg/mL Tricaine (MS222/ethyl 3-aminobenzoate; Sigma-Aldrich) until fish became non-responsive to touch (<5 minutes). In some cases, additional tricaine was added dropwise to achieve timely non-responsiveness. Anesthetized fish were moved to an imaging chamber with a transfer pipet (Fisher 13-711-7M) cut halfway to the base to increase the hole size. Depending on the size of the specimen, fish were either maintained in 4-well (Ibidi 80427), 2-well (Ibidi 80287), or 1-well (Nunc Lab-Tek II 155360) glass-bottom imaging chambers. The smallest possible chamber was always used and enough water with tricaine was added to just cover the fish body. Drift was limited by adding soaked sponge cloth to the chamber around the fish, and additional soaked sponge cloth was draped over the fish head and upward facing gills to maintain viability. In all cases, fish were imaged for 20 minutes. Following imaging, fish were kept in 0.16 mg/mL Tricaine for 20 minutes to ensure euthanasia. For non-terminal experiments, fish were quickly transferred to a tank of fresh water RO water to promote recovery following imaging experiment. We found fish could tolerate <10 minutes in 0.08-0.16 Tricaine and reliably recover.

For whole-fish confocal tiling images, the fish were first completely euthanized and then immobilized in a 1-well glass-bottom chamber (Nunc Lab-Tek II 155360) with low gelling temperature agarose (Sigma Aldrich 19045). To visualize scales, we immersed euthanized fish into 0.04% solution of Alizarin red powder (Sigma Aldrich 130-22-3) dissolved in RO water in a 6-well plat for 20 minutes. Fish were then washed for three times for 5 minutes in fresh RO water with gentle rocking prior to imaging.

### Puncture Wound and Listeria Infection Model

A streak plate from L. monocytogenes strain 10403S frozen stock was grown at 37°C. A fresh colony was picked and grown statically in 1 mL brain–heart infusion (BHI) medium overnight at 30°C to reach stationary phase. Bacteria were sub-cultured for ∼1.5-2 hr in fresh BHI (4:1, BHI:overnight culture) to achieve growth to mid-logarithmic phase (OD600 ≈ 0.6–0.8). 3 mL of the mid-logarithmic phase bacterial culture were spun down and washed three times in phosphate-buffered saline (PBS; Gibco 14190-144) and resuspended in 100 µL of PBS for infection. This bacterial suspension was then added to 10 mL of water in a 6-well plate. Adult zebrafish were anesthetized in 0.16 mg/mL Tricaine and transferred to a damp sponge cloth. A 30G½ needle (Becton Dickinson PrecisionGlide) was used to puncture the caudal peduncle on the dorsal side of the animal. Pressure was applied until the needle came out the other side. Wounded fish were placed in a tank of water until movement was observed (0-3 minutes) at which point they were added to the 6-well plate with bacteria. This plate was gently rocked for 2 hours in the dark at which point the fish were moved to a 1 liter tank with fresh water and maintained in a laboratory 28.5°C incubator until ready for imaging. Water was exchanged daily and fish were fed 1-2 times daily.

### Microscopy and Image Preparation

For imaging the TLN over development (Figure 1B), fish were imaged on a Zeiss Zoomscope (EMS3/SyCoP3; 1× Plan-NeoFluar Z objective; Zeiss) with an Axiocam Mrm charge-coupled device camera using ZenPro 2012 software (Zeiss). For most other fish imaging, we used a spinning disc confocal microscope CSU-X, Yokogawa, Sugar Land, TX) with a confocal scanhead on a Zeiss Observer Z.1 inverted microscope, either an EC Plan-Neofluar 10x/0.30 or EX Plan-Neofluar 40x/0.75 object, a Photometrics Evolve EMCCD camera and Zen software (Zeiss). To stitch tiled images in ZenPro 2012 software (Zeiss), we first performed a max intensity *z*-projection and then stitched with a 10% maximal shift. All images were prepared for publication with ImageJ (v. 2.1.0/1.53c)

### Cell Tracking

Cell movies to be tracked were imported into ImageJ (v. 2.1.0/1.53c), and individual cells were tracked using the Manual Tracking plugin. Some movies were rotated prior to tracking to ensure that dorsal side of the fish was always pointing straight upward. Even when the CD4:mCherry channel is not show, all cell tracking was performed using dual labelled *tg(lck:GFP/cd4-1:mCherry)* fish as differentiating and tracking individual T cells was prohibitively difficult otherwise.

For Figure 3, every trackable CD4+Lck+ cell was tracked. Cells were only excluded if they left the field of view during the imaging window. CD4-Lck+ cells appeared to move indistinguishably from CD4+Lck+ cells, although we did not attempt to track them. For Figure 4, the cell density around the infected wounds was too high to track every cell. Instead, we tracked 20 randomly chosen (CD4+ or CD4-) motile T cells from every video. Cell tracking XY coordinates were copied into Microsoft Excel (v. 16.65), which was used to calculate velocity, directionality, displacement, and mean squared displacement. Additionally, Excel was used to generate the cell track rose plots, which were then overlaid in Illustrator (Adobe Inc.; v. 26.02) with graphs of the fish-by-fish cell averages to generate complete figures.

### Dissection and cell isolation from the gut, gills, spleen, and kidney

We dissected adult zebrafish as previously described (47). Briefly, we used dissecting scissors to cut along the belly of euthanized adult zebrafish to expose the internal organs. The spleen was separated from the gut with curved tweezers and added to a 40-micron filter in a 6-well plate containing 8 mL of cell isolation media (L-15 + 1 mM EDTA, penicillin-streptomycin, and 2% FBS). The gut was then added to a 1.5 mL Eppendorf tubed filled with 1 mL pre-warmed 37°C collagenase/dispase solution (Roche 10 269 638 001) and incubated for 90 seconds before adding to a 40-micron filter as above. The operculum, pectoral fin, and pectoral girdle were then all cut out, and the intact gills were removed from with curved tweezers and added to a dish of PBS. Non-gill tissue was then carefully removed under a dissecting microscope. In 3 of 6 fish, only half the gill arches were cleanly isolated, so final cell counts were doubled in these cases to get estimates of the gill totals. Finally, curved tweezers were used to isolate the entirety of the kidney, which was then added to a 40-micron filter in a 6-well plate filled with 8 mL of cell isolation buffer. All organs were then mechanically disrupted with the back end of a plastic syringe, and the resulting filtered cell suspension was used for further analysis.

### Descaling, TLN cell isolation, and flow cytometry

Euthanized adult zebrafish were briefly dunked in room temperature PBS and then placed in a 60mm Petri Dish (Fisher FB0875713A) containing 8 mL of room temperature PBS. One (gloved) hand was used to hold the fish in place by firmly grasping the head, and the second hand was used to scrape rostrally along the body of the fish with an angled dissecting knife (Fine Science Tools 10056-12) to descale. This was done under a dissecting microscope and scale loss was visually monitored. Scales at the base of the caudal, dorsal, and anal fins typically had to be individually removed by plucking with thin-tip tweezers (Dumont). PBS from the Petri Dish was then washed over the scaled fish for approximately 30 seconds with a transfer pipette (Fisher 13-711-7M) to promote cells entering the suspension, and the buffer was then passed through a 40-micron filter. The pressure from descaling often caused males and females to release their eggs and milt, respectively. For this reason, only females were used for analysis, as their eggs could be readily filtered out. This cell suspension was then mixed 1:1 with cell isolation media in a 15 mL falcon tube, washed, resuspended in 500 µL of ACK lysis buffer for 90 seconds to lyse red blood cells, and washed once more with cell isolation media. For Figure 2C, these cells were added to an 8-well chamber glass-bottom chamber (Ibidi 80807) and imaged at room temperature on the Zeiss Zoomscope (EMS3/SyCoP3; 1× Plan-NeoFluar Z objective; Zeiss) with an Axiocam Mrm charge-coupled device camera using ZenPro 2012 software (Zeiss). For flow cytometry, cells isolated from tg(Lck:GFP/cd4.1:mCherry) by descaling were stained with Ghost Dye™ Red 780 (Tonbo) per manufacturer’s instructions, washed with twice with FACS buffer (PBS + 1 mM EDTA + 5% FBS + penicillin/streptomycin), and analyzed by flow cytometry. A ThermoFisher Attune was used for all experiments, allowing quantitation of total cell numbers of interest in the sample.

### Single Cell RNA-sequencing

For scRNA-seq, cells were isolated from uninfected control and 4 days post-infection zebrafish exactly as described above except that all buffers were prepared without EDTA. To enrich for cells involved in the immune response, we only descaled the caudal peduncle rather than the entire fish to harvest cells for scRNA-seq. Libraries were constructed from these cells by the University of Wisconsin-Madison Biotechnology & Gene Expression Center (research resource identifier [RRID]: SCR_017757) using the Chromium Single Cell Gene Expression Solution 3’ v2 (10x Genomics) and sequenced by the DNA Sequencing Facility (RRID: SCR_017759) using NovaSeq6000 (Illumina) with read lengths of 29-bp + 90-bp (Read1 + Read2). Raw reads were processed by *cellranger count* (v6.1.2, 10x Genomics) with default parameters for read tagging, alignment to zebrafish reference genome (GRCz11), and feature counting based on an improved zebrafish transcriptome annotation (48). In total, 386M and 486M reads were sequenced for the Control and Infected samples respectively, both with valid barcode rates over 95% and a valid UMI rate of 99.9%.

### Unsupervised clustering and cell type identification

Filtered cell-by-gene count matrices generated were used for graph-based clustering with Seurat v4.1.1 (49). We filtered out genes that are detected in less than four cells and cells with less than 200 genes detected or a mitochondrial ratio > 5%. Doublets were also removed using DoubletFinder with an estimated rate of 3% (50). Post doublet removal, the Control sample has 3141 cells with a median of 1271 captured genes, while the Infected sample has 4309 cells with a median of 2125 captured genes.

Normalization (SCTransform), and dimensional reduction were done with default parameters in Seurat package prior to clustering and UMAP projection (51). Clustering and visualizations were either performed on the Control sample only, or with both samples integrated via canonical correlation analysis (CCA)-based method from Seurat (52). To identify cell types and states at chosen clustering resolution (0.6 for Control only, 0.5 for Control and Infected integrated), we performed differential expression analysis using Wilcoxon rank sum-based method (log2 fold change > 0.25, % of expressing cells > 0.25). Known markers were used to identify T cells (*cd247l, lck, il7r*, and *cxcr4a*) (53-55), dendritic cell-like populations (*ctsbb, tlr7*) (56, 57), B cells (*cd37, pax5*) (58, 59), macrophages (*mpeg1*.*1, grn1*) (60, 61), neutrophils (*mpx, il6r*) (11, 62), erythrocytes (*hbba2, hemgn*) (63), thrombocytes (*thbs1b, fn1b*) (57), superficial epithelial cells (*krt1-19d, cldne*) (64), ionocytes (*trpv6, foxi3b*) (65), intermediate epithelial cells (*cldna, tp63*) (64), basal epithelial cells (*cldn1, cldni*) (64), mesenchymal cells (*vcana, clu*) (66), lateral line-like cells (*prox1a, prr15la*) (67), and epidermal mucous cells (*agr2, cldnh*) (64). We found one population with some cells locating closer to the T cell populations but without clear enrichment for known signatures and included this population for a subclustering analysis with other T cells. We identified one subcluster with epithelial features but without known T cell features, thus labeled them as Others-epithelial cells and removed from the T cell-focused analysis. T cell subsets were identified using known marker gene expression patterns: naïve T cells (*ccr7, cxcr4a*) (68), cycling T cells (*mki67, top2a*) (69), regulatory-like T cells (*foxp3a, ccr6a, ccl20a*.*3*) (70, 71), cytotoxic T cells (*cxcr3*.*1, gzma, prf1*.*1, nkl*.*2, ccr2, prf1*.*9*) (72), *ccl38*.6-high T cells (*il2rb, tnfsf14, ccl38*.*6*), Th2 cell/ILC2 (*il11b, il4, il13*), *rag1/2*+ T cells (*bcl11ba, rag1, rag2*) (57), lymphocyte-like cells (*ccl33*.*3, lcp2b*). For a more stringent T cell-focused analysis in the Control and Infected integrated sample, we extracted T cells using the expression of both cd247l and lck (log2FC > 0.3, % of expressing cells > 0.3).

### Statistical Analysis

Statistical tests were all performed using GraphPad Prism (version 9.4.1), with specific tests indicated in each figure legend. P values were denoted on graphs as follows: n.s., P > 0.05; *, P < 0.05; **, P < 0.01; ***, P < 0.001; ****, P < 0.0001.

## Acknowledgements

We would like to thank A. Horn, V. Miskolci, J. Squirrell, D. Bennin, T. Schoen, M. Giese, A. Fister, G. Ramakrishnan, A. Peterson, and N. Mercado-Soto for their helpful discussion throughout the design and execution of this project. We would also like to thank the Gene Expression Center at the University of Wisconsin-Madison for sequencing assistance, and the Carbone Center Flow lab for assistance in training and guidance of flow cytometry experiments. Figures throughout this paper were made in part with biorender.com.

## Funding

National Institute of Health grant 1 F32 GM146398-01 (T.F.R)

National Institute of Health grant T32 HL07899 (T.F.R.)

National Institute of Health grant R35 GM118027 (A.H.)

## Data and materials availability

Most data are available in the main text or the supplementary materials. Images beyond the representative ones shown in this report, raw cell tracks, and subsequent analysis are all available upon request. scRNA-seq data are available in the NCBI Expression Omnibus (GSE215189) and is available to reviews of this manuscript with the token: azgjgwmwpzknvkr. Upon publication, this data will be made publically available. scRNA-seq analysis codes are available at: https://github.com/k326xh/TLN-Tcell-scRNA-analysis.

## Supporting Information for

### Supplemental Figures and Legends

**Supplemental Figure 1.**
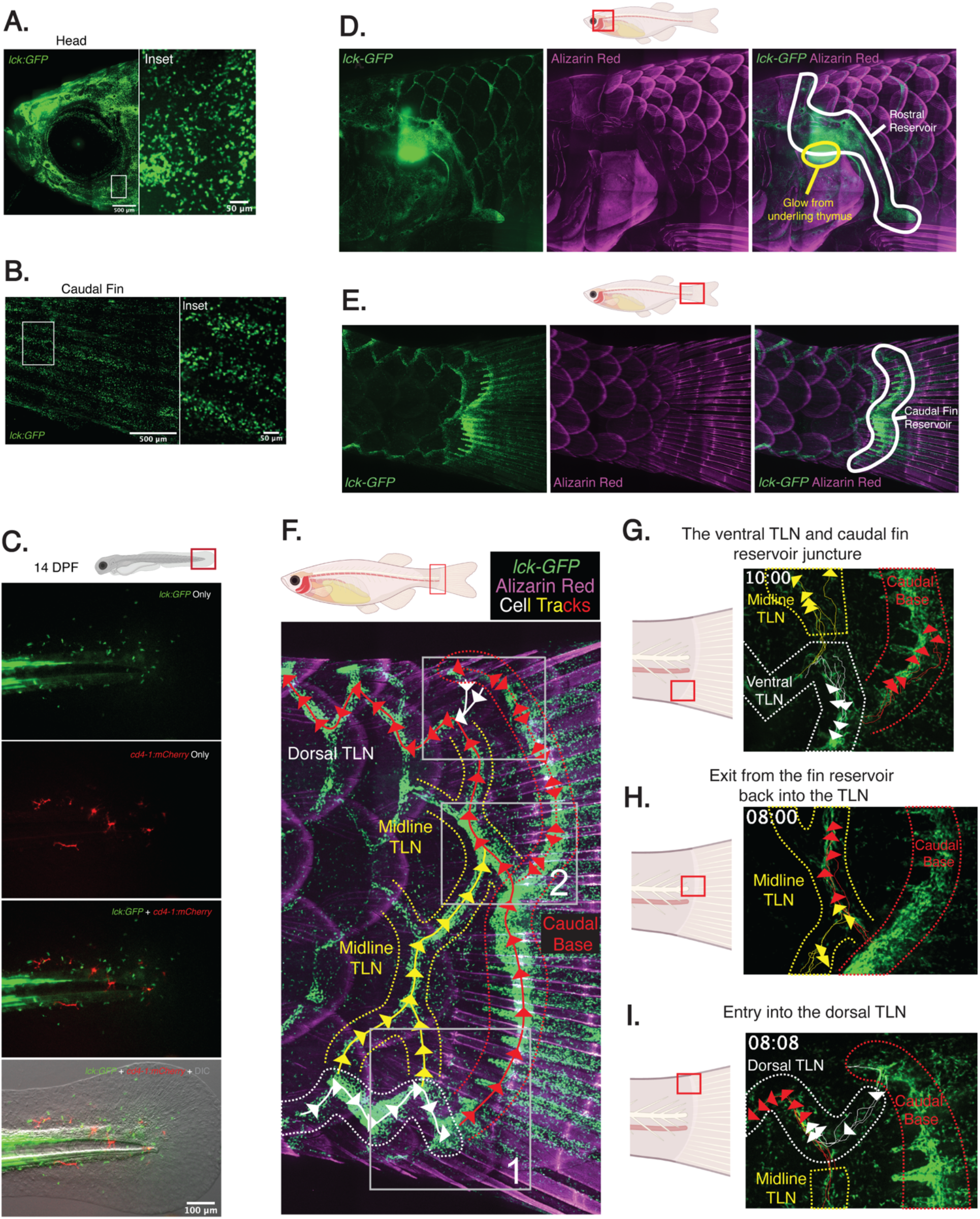
Characterization of T cells and their movement within the TLN. **(A-B)** Portions of the whole-body confocal tiled image of a 10 wpf lck:GFP fish from Figure 1A cropped to show the head (A) and caudal fin (B). Insets show the diffuse organization of T cells in these regions. Note that the striped pattern observed in the fins (B) is due to T cells primarily occupying the space between the bony rays. **(C)** A representative image of the caudal fin region of a 14 dpf transgenic lck:GFP;cd4-1:mCherry larvae shows the presence of mostly mCherry-negative T cells as well as GFP-mCherry+ macrophages. **(D-E)** An image of the rostral (D) or caudal (E) end of the body of an lck:GFP zebrafish stained with Alizarin red shows the abundance of GFP-expressing cells that make up the rostral and caudal reservoirs. **(F)** A still image of the caudal fin reservoir and surrounding TLN from a transgenic lck:GFP zebrafish stained with Alizarin red. Three numbered boxes have been overlaid corresponding to the regions imaged in (G-I). Color-coded cell tracks are overlaid on the image corresponding to the results from (G-I). **(G)** Color-coded cell tracks illustrate the three basic T cell behaviors at the juncture between the ventral TLN and the caudal fin reservoir, corresponding to position 1 in (F). Yellow tracks show some T cells migrating dorsally even before reaching the reservoir. White tracks show T cells entering the reservoir from the ventral TLN. Red tracks show T cells already in the caudal base displaying ventral to dorsal movement towards the midline of the fish. **(H)** Color-coded cell tracks illustrate two migration behaviors near the midline of the fish, corresponding to position 2 in (F). First, yellow tracks show T cells not within the reservoir migrating dorsally. These cells had likely left the ventral TLN before reaching the reservoir, matching the behavior of the yellow tracks in (F). Second, red tracks show T cells exiting the caudal fin reservoir at a distinct site just dorsal to the midline, re-entering the TLN, and exhibiting coordinated ventral to dorsal movement. **(I)** Color-coded cell tracks show cell migration behavior at the juncture between the caudal fin reservoir and the dorsal TLN, corresponding to position 3 in (F). This still image is taken from Movie S6. Red cell tracks show T cells in the midline TLN moving dorsally and ultimately reentering the dorsal TLN, resuming their zigzagging motility now in the rostral direction. White tracks show a small number of T cells exiting the caudal base reservoir at the dorsal-most point and directly entering the dorsal TLN. Note that for (G-I) representative tracks illustrating the overall behavior rather than every single cell is shown.

**Supplemental Figure 2.**
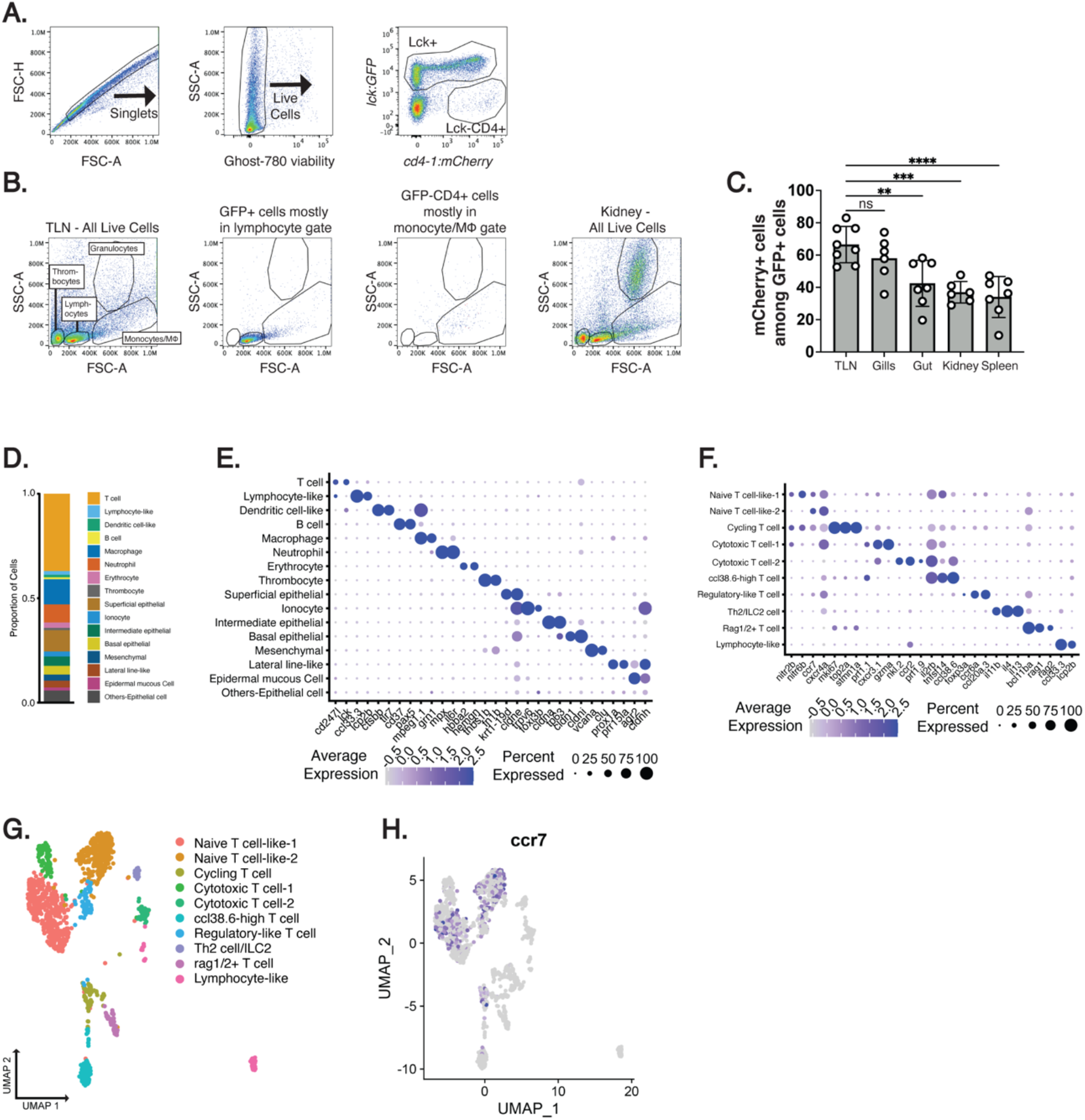
Analysis of the TLN by flow cytometry and scRNA-seq. **(A)** Flow cytometry plots show the basic gating strategy used for quantifying cell populations from the double transgenic *lck:GPF;cd4-1:mCherry* zebrafish. **(B)** Flow cytometry plots show that the isolated GFP-expressing cells localize to the expected lymphocyte gate and that GFP-negative, mCherry-positive cells mostly localize to the monocyte/macrophage gate. A plot from the granulocyte-rich kidney is also shown to demonstrate how the granulocyte gate was drawn. **(C)** A graph shows the percentage of GFP-expressing cells that also express mCherry from double transgenic *lck:GFP;cd4-1:mCherry*. Each dot corresponds to an individual fish pooled from three independent experiments. The mean percentage of the TLN was compared to the mean percentage from all other sites using an ordinary one-way ANOVA. **(D)** Proportions of all cell types in the TLN determined by scRNA-seq. **(E)** Representative gene expression in each of the major cell types on dot plot. Color gradient: normalized relative expression level. Dot size: percentage of cells in the cluster that express the specified gene. **(F-G)** Subclustering of T cells reveals two major naïve-like T cell populations, with gene expression data shown as a dot plot (F) or as UMAP analysis (G). **(H)** An overlay of *ccr7* expression of the UMAP analysis from (G) shows that *crr7* expression is mostly found in the two naïve-like populations.

**Supplemental Figure 3.**
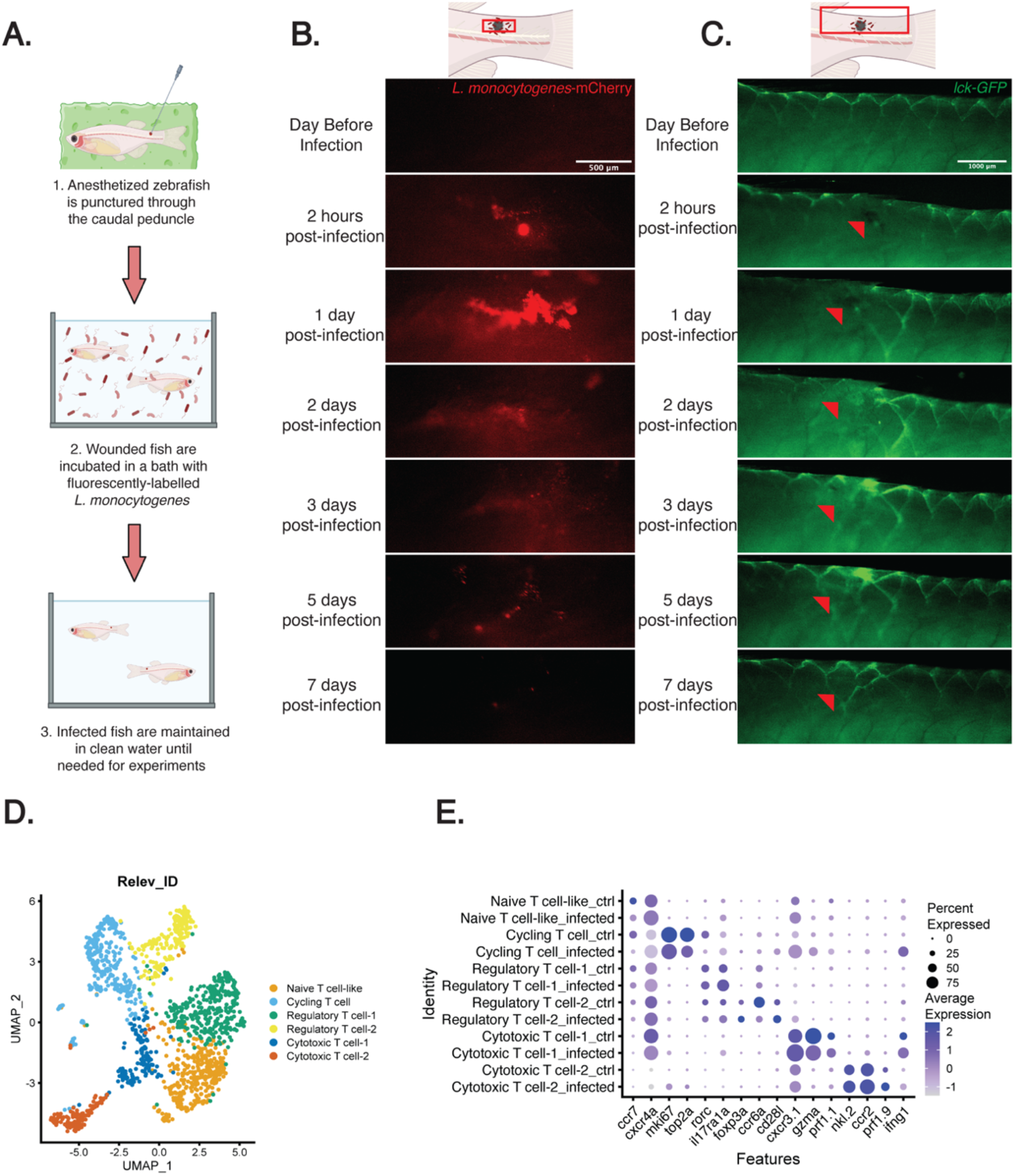
Analysis of TLN T cells in control and *listeria-*infected fish. **(A)** A cartoon illustrates the puncture wound infection technique with accompanying descriptions. **(B)** A series of high-magnification images show fluorescence signal from the mCherry-expressing *L. monocytogenes* near the puncture wound of the same fish over a 7-day time course. For the day before the image, we imaged the approximate location of the future puncture. **(C)** A series of images show how the TLN changes in the same *lck:GFP* fish following puncture wound infection over a 7-day time course. The red arrow indicates the site of the puncture wound, which is apparent up to 1-day post-puncture and approximated thereafter. **(D)** UMAP illustration of TLN T cells colored by subtypes. **(E)** Representative gene expression in T cell subtypes by conditions shown on dot plot. Color gradient: normalized relative expression level. Dot size: percentage of cells in the cluster that express the specified gene.

### Supplemental Movie Legends

**Movie S1**.

Live confocal imaging of the TLN of an adult *tg(lck:GFP;cd4-1:mCherry)* shows the characteristic streaming motility with a 10x objective to highlight T cell movement through the TLN network over hundreds of microns. This movie corresponds to Figure 1F, where individual cell tracks are shown. Time in mm:ss.

**Movie S2**.

Live confocal imaging of the TLN of an adult *tg(lck:GFP;cd4-1:mCherry)* shows the directional movement of T cells. Three representative cells are tracked (white dot and lines) corresponding to the outlined cells in Figure 3A. Time in mm:ss.

**Movie S3**

Live confocal imaging of the TLN of an adult *tg(lck:GFP;mpeg1::mCherry*^*CAAX*^) display the streaming motility behavior of T cells. Two side-by-side representative movies are show, with the left movie corresponding to figure 3I. Macrophages are distributed throughout the network. Individual T cells are difficult to track by just the GFP channel, but the movie demonstrates T cells streaming past TLN-localized macrophages under homeostatic conditions. Time in mm:ss.

**Movie S4**.

The left portion of this video shows a still image of a region proximal to the puncture wound *l. monocytogenes* infection of a transgenic *lck:GFP* adult zebrafish. The two boxed regions in the still image are shown as movies on the right. The ‘motile’ region shows a group T cells that mostly migratory, and the ‘sessile’ region shows a group that is mostly non-migratory. Time in mm:ss.

**Movie S5**.

Live imaging of the region proximal to the puncture wound *L. monocytogenes* infection of an adult transgenic *lck:GFP;cd4-1:mCherry* zebrafish. The video first shows still frames with the relevant macrophages labelled (M) and a white dot over the GFP+mCherry+ T cells. Once the movie starts, the white dot follows the T cell as it contacts all three of these infection-proximal macrophages. Time in mm:ss.

## Notes

### Competing Interest Statement

The authors have declared no competing interest.

